# Synthetic MPRA perturbation supervision improves variant-effect prediction in sequence-to-function models

**DOI:** 10.64898/2026.11.02.763699

**Authors:** Lena Wang, Jake Galvin, Eduarda Vaz, Rebecca Keener, Alexis Battle

## Abstract

Deep learning sequence-to-function (S2F) models learn to predict functional readouts, such as gene expression, from genetic sequence. However, these models are trained predominantly on the reference genome and struggle to accurately predict the effect of genetic variation on gene expression. Massively parallel reporter assays (MPRAs) explore the direct impact of genetic variation on gene expression, but saturation-mutagenesis MPRAs that measure variant effects directly remain limited, thereby restricting their use for fine-tuning S2F models. In this work, we investigate fine-tuning Borzoi, a state-of-the-art S2F model, on model-generated synthetic MPRA variant-effect data as a scalable alternative to overcome limits on the availability of supervised training data for genetic variation. First, we trained an ensemble of 16 models on 110,359 K562 MPRA measurements. Then, we distilled their predictions into a single student model using MPRA elements associated with 1,000 genes. The distilled model was subsequently used to generate synthetic variant-effect labels across increasing numbers of genes (50 to 1,000), with 600 variants per gene, to fine-tune Borzoi. Performance improved as we included more genes in training. In data-limited settings, training on more genes with fewer variants per gene performed better than training on fewer genes with more variants per gene, given the same number of total variants. Compared to pre-trained Borzoi, Borzoi fine-tuned on 600,000 synthetic samples across 1,000 genes increased Pearson correlation from 0.28 to 0.54 on CAGI5 promoter variants and from 0.39 to 0.46 on 903 GTEx fine-mapped whole-blood expression quantitative trait loci (eQTLs). Importantly, these improvements were achieved while preserving the pre-trained performance across all 7,611 output tracks. Furthermore, fine-tuned Borzoi recovered regulatory motifs identified by the MPRA model in genes unseen during training, suggesting that learned regulatory features can generalize across loci. Together, these results provide a proof of concept that model-generated synthetic supervision can improve variant effect prediction in S2F models while preserving their original pre-trained capabilities, offering a scalable strategy toward capturing the impact of genetic variation on gene regulation.

## Introduction

Deep learning sequence-to-function (S2F) models predict molecular phenotypes from genetic sequence, learning robust correlations between genomic features and their corresponding functional readout^1–3^. This pairing yields strong predictive performance and enables generalization to unseen loci. Yet as S2F models have grown more complex and come to include more functional data, their underlying training paradigm has remained unchanged, using primarily the reference genome without exposure to the effects of sequence perturbation. The latest S2F models still struggle to accurately predict the effect of *cis*-regulatory genetic variation on gene expression, limiting their utility in downstream applications^4–6^.

One approach to address this performance gap is to fine-tune S2F models directly on personal genomes with paired expression data, achieving performance on par with established linear methods on genes included in the training data across individuals. However, models fine-tuned on personal genomes appear to generalize poorly to genes and variants not seen during training, relying heavily on memorizing the effects of strong, common variants such as eQTL variants^7–9^ to drive their performance among previously seen genes. We have shown that although larger training sets including personal genomes improve prediction accuracy somewhat^10^, there is still room for greater improvements. These limitations highlight the need for more diverse variant-level supervision. However, experimentally measured regulatory variant effects are costly and remain available for only a small fraction of genetic variants, limiting their usage for fine-tuning S2F models^11^. Another challenge in adapting S2F models to new tasks is preserving their pretrained predictive capabilities. Existing fine-tuning approaches often replace pre-trained genomic output tracks with task-specific prediction heads, reducing the fine-tuned model’s performance on the original prediction tasks^7,8,10^. Recent work demonstrated that replaying pretraining data when fine-tuning on synthetic perturbation can maintain pre-trained task performance while improving generalization to unseen perturbations^12^. However, this approach requires access to the original pretraining data, motivating alternative approaches that preserve pretrained capabilities without relying on data replay.

We address these limitations by fine-tuning Borzoi^3^, a state-of-the-art S2F architecture, using only synthetic variant effect size labels while preserving Borzoi’s original output heads. We generated large-scale synthetic variant-effect training data from an ensemble of small, specialized models trained on massively parallel reporter assay (MPRA) data^13,14^. This provides a general framework for transferring regulatory knowledge learned from specialized models to large S2F models. We show that this approach improves variant effect prediction, including for enhancer variants and genes unseen during training, while preserving Borzoi’s original capabilities.

## Results

### Overview: Fine-tuning Borzoi on synthetic MPRA data improves variant-effect prediction

We developed a framework to fine-tune Borzoi on synthetic variant effect labels to improve its prediction and generalization of how genetic variants affect gene expression (Fig. 1a). We selected 1,000 genes spanning a range of *cis*-heritability (0.01–0.78), estimated based on *R*^*2*^ performance of elastic net predictions on held-out test individuals from GTEx wholeblood^15^. For each gene, we performed *in silico* mutagenesis of the 200-bp core promoter, as annotated in an existing MPRA dataset^13^, yielding 600 SNVs per gene and 600,000 SNVs in total. We first constructed a distilled MPRA model and used it to generate synthetic labels for each of the 600,000 SNVs by calculating the difference in predicted activity between 281-bp reporter constructs carrying the reference or alternative alleles (detailed in the next section). We used these synthetic labels, along with predicted expression coverage of the reference sequence from pre-trained Borzoi, to fine-tune Borzoi on matched reference and alternative 49,152-bp sequences centered on each variant. We used a pairwise loss function that aligns Borzoi’s predicted variant effect with the synthetic label, where the fine-tuned Borzoi model predicts its effect as the log_2_ fold change in predicted whole-blood RNA-seq coverage summed over the target gene’s exons, thereby contextualizing variant effect predictions within the model’s existing outputs. However, unconstrained fine-tuning risks disrupting predictive capabilities acquired during pre-training. We therefore regularized reference-sequence predictions by incorporating a Poisson loss^3^ between the predicted functional coverage from pretrained and fine-tuned Borzoi on the reference sequence across all 7,611 output tracks.

**Figure 1:**
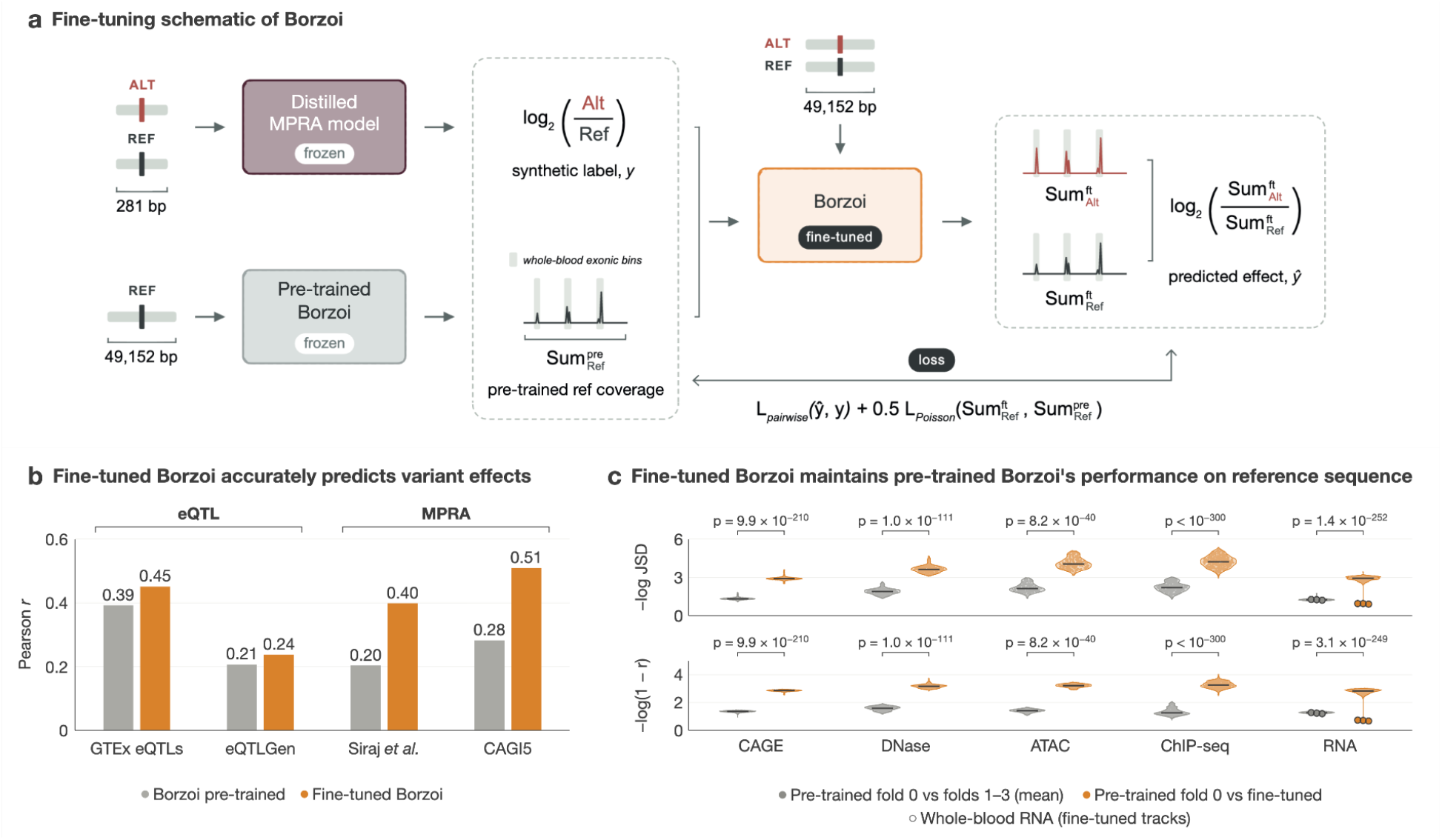
Fine-tuning Borzoi on synthetic MPRA data improves variant-effect prediction. **a**, Fine-tuning scheme of Borzoi. A pairwise loss fits the distilled MPRA student’s predicted effect for each variant *y* to Borzoi’s predicted variant effect ŷ. A Poisson term keeps the fine-tuned model aligned with the frozen pre-trained model. **b**, Variant-effect prediction on GTEx eQTLs (*n* = 903), eQTL Gen (*n* = 3,539), Siraj *et al*. K562 MPRA activity (*n* = 14,172), and CAGI5 saturation mutagenesis in eight promoter elements (*n* = 5,769). Differences between pre-trained and fine-tuned Borzoi were tested with gene-level bootstrap for GTEx (*p* = 0.01), eQTLGen (*p* = 0.095), and Siraj et al. (*p* < 0.001). A two-sided paired Wilcoxon test was used for CAGI5 (*p* = 0.0078). **c**, Predictions from pre-trained fold 0 were compared with three other pre-trained replicates (*grey*) and fine-tuned Borzoi (*orange*) across 1,436 genes and all 7,611 human tracks. Deviations were quantified by Jensen-Shannon divergence (*top*) and Pearson correlation (*bottom*). Statistical significance was assessed using paired Wilcoxon tests.

We first evaluated fine-tuned Borzoi against its pre-trained model on four experimental variant effect prediction benchmarks (Fig. 1b). On 903 fine-mapped GTEx whole-blood eQTLs (PIP > 0.9) in genes held out from training, fine-tuning improved Pearson *r* from 0.39 to 0.45. On 3,539 eQTLGen whole-blood eQTLs, it improved *r* from 0.21 to 0.24. On the MPRA saturation mutagenesis benchmarks, fine-tuning improved *r* from 0.20 to 0.40 on the Siraj *et al*. dataset^16^ and from 0.28 to 0.51 on CAGI5. Consistent with our objective of preserving predictive capabilities of pre-trained Borzoi, fine-tuned Borzoi model predictions remained closer to its starting checkpoint (pre-trained Borzoi replicate 0) than those of independently pretrained replicates, as measured by Jensen–Shannon divergence (JSD; -log_10_(JSD) = 3.72 versus 1.86) and Pearson *r* (-log_10_(1 - *r*) = 3.07 versus 1.31) across all 7,611 output tracks, including CAGE, DNase-seq, ATAC-seq, ChIP-seq, and RNA-seq (Fig. 1c). Thus, fine-tuning on synthetic MPRA labels improves Borzoi’s variant effect predictions while preserving its pre-trained predictions on reference sequences.

### Distillation yields a robust MPRA model to generate synthetic variant-effect labels

To generate synthetic labels for supervision, we built an initial MPRA model by fine-tuning AlphaGenome’s pre-trained encoder on K562 lentiMPRA elements from Agarwal et al., following the protocol inspired by Murphy et al. Ensembling can improve the accuracy and out-of-distribution generalization of genomic models, and distillation can retain these gains in a single model at lower inference cost^17^. We therefore trained an ensemble of MPRA teachers with chromosome-level cross-validation and distilled it into a single model of the same architecture (Fig. 2a). To generate distillation data, we performed *in silico* mutagenesis of the 200-bp Agarwal et al. promoter and enhancer elements mapping to the 1,000 training genes. Each teacher predicted reference and alternative allele activity with the variant at four positions (centered and three offsets within ±20 bp), in both forward and reverse-complement orientations. We averaged these eight predictions within each teacher, then across teachers, and trained the distilled model to reproduce the ensemble mean for each allele. Models distilled from eight teachers outperformed those distilled from four (*r* = 0.59 versus *r* = 0.58, *p* < 10^−4^) and performed comparably to those distilled from 16 (Supplementary Fig. 1). We used the model distilled from 16 teachers for subsequent analyses to maximize ensemble diversity.

**Figure 2:**
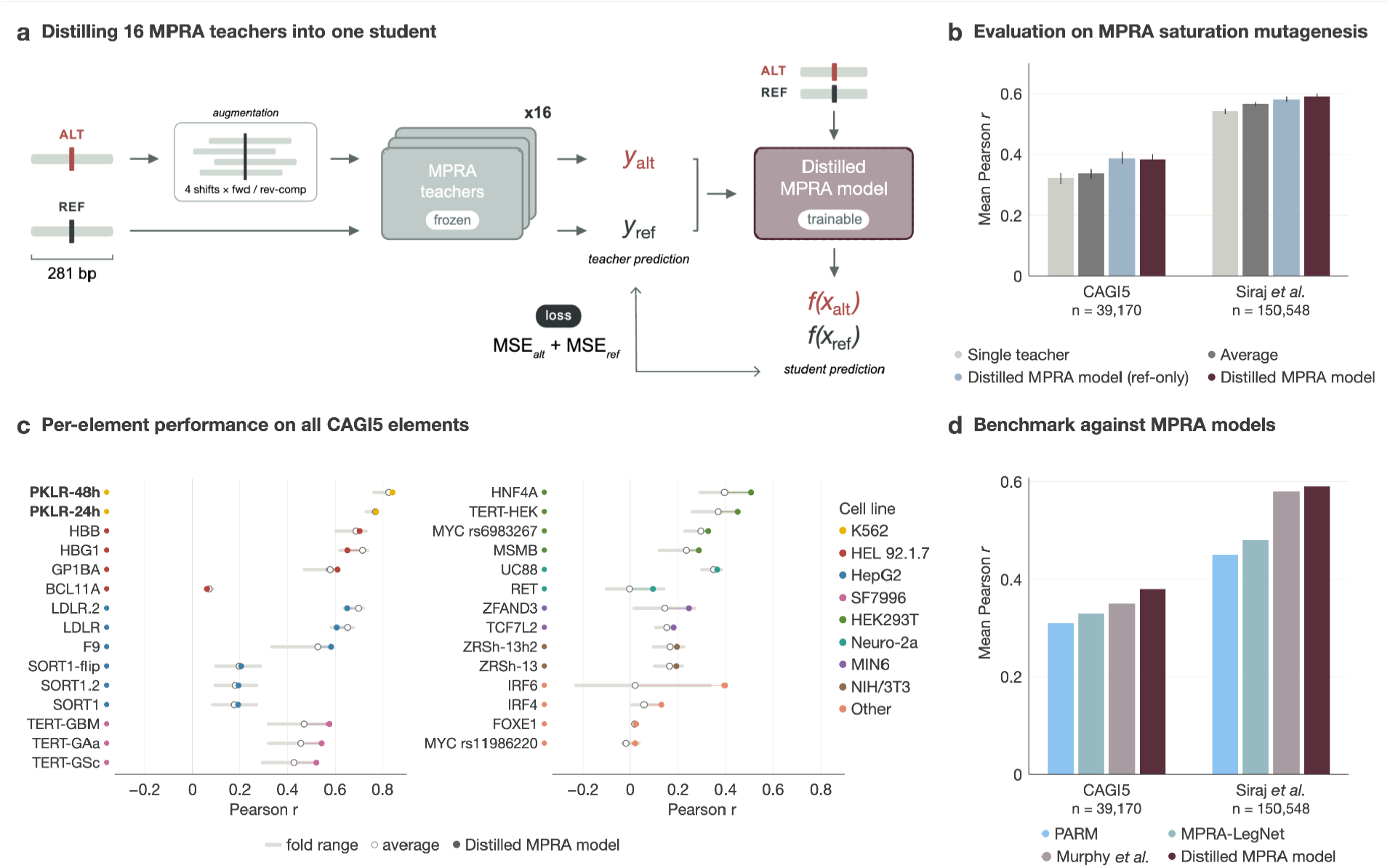
Ensemble distillation yields a robust MPRA model. **a**, Schematic for MPRA model distillation. For each synthetic variant, each teacher makes a reference- and alternative-sequence prediction using four positional offsets in both forward and reverse-complement orientations. The distilled student model is trained to reproduce ensemble-averaged predictions. **b**, Mean Pearson *r* on CAGI5 (29 elements; *n* = 39,170) and Siraj *et al*. K562 MPRA activity (102 elements; *n* = 150,548) for individual teacher folds, the 16-fold mean, student model distilled from reference-allele predictions, and student model distilled from reference- and alternative-allele predictions. **c**, Per-locus Pearson *r* on CAGI5. Grey bars range across 16 individual teachers. Open circles represent the average of 16 teachers. Filled circles represent fully distilled MPRA students. **d**, Comparison of variant-effect prediction performance between the distilled MPRA student, AlphaGenome-based MPRA activity model trained by Murphy *et al*., MPRA-LegNet, and PARM on CAGI5 (*left*) and Siraj *et al*. (*right*). Performance was measured by mean per-element Pearson correlation.

We evaluated the distilled model on two MPRA saturation mutagenesis datasets: CAGI5 (29 regulatory elements across 12 cell lines) and Siraj *et al*. (102 elements in K562). We compared individual teachers, the 16-teacher ensemble average, and models distilled from reference predictions alone or from both reference and alternative alleles (Fig. 2b). Ensembling increased the mean Pearson *r* over individual teachers (CAGI5: 0.32 to 0.34; Siraj et al.: 0.54 to 0.57), and distillation improved accuracy further (0.38 and 0.59). Distilling from both alleles matched reference-only distillation on CAGI5 and outperformed it on Siraj et al. On CAGI5, the distilled model (0.38) also outperformed existing MPRA models, including PARM^18^ (0.31), MPRA-LegNet^13^ (0.33), and the AlphaGenome-based model of Murphy *et al*.^14^ (0.35), which shares its architecture but lacks ensemble distillation (Fig. 2d). Across CAGI5 elements, the distilled model matched or exceeded the ensemble average for 26 of 29 elements (Fig. 2c), including 23 of 27 elements assayed outside K562, suggesting that the benefits of distillation extend beyond the training cell type. The highest prediction accuracy was observed for *PKLR*, assayed in K562 cells. Together, these results show that ensemble distillation yields an accurate MPRA model for generating synthetic labels to fine-tune Borzoi.

### Fine-tuning gains depend on context, scale, and regulatory signal

Given fine-tuning was restricted to promoter variants, we first asked whether the gains were restricted to local promoter variants or extended to broader genomic settings. We evaluated GTEx whole-blood eQTLs overlapping promoter-like or enhancer-like ENCODE candidate *cis*-regulatory elements (cCREs) (Fig. 3a). Fine-tuning increased Pearson *r* from 0.50 to 0.56 for 169 promoter-like variants and from 0.48 to 0.59 for 327 enhancer-like variants. We also compared against PromoterAI^19^, which was fine-tuned on approximately 58,000 rare promoter variants with expression effects estimated from RNA-seq measurements. Despite using only synthetic variant-effect labels during Borzoi fine-tuning, our model achieved similar performance to PromoterAI on variants in promoter-like elements (*r* = 0.56 versus *r* = 0.58) and higher accuracy on variants in enhancer-like elements (*r* = 0.59 versus *r* = 0.51). Fine-tuned Borzoi also exceeded the distilled MPRA model in both promoter-like and enhancer-like variants (*r* = 0.56 versus *r* = 0.47 and *r* = 0.59 versus *r* = 0.49, respectively). These results emphasize that the fine-tuned model retains broader knowledge about genomic regulatory patterns than what is provided by the synthetic MPRA dataset and that this approach can improve predictions beyond promoters.

**Figure 3:**
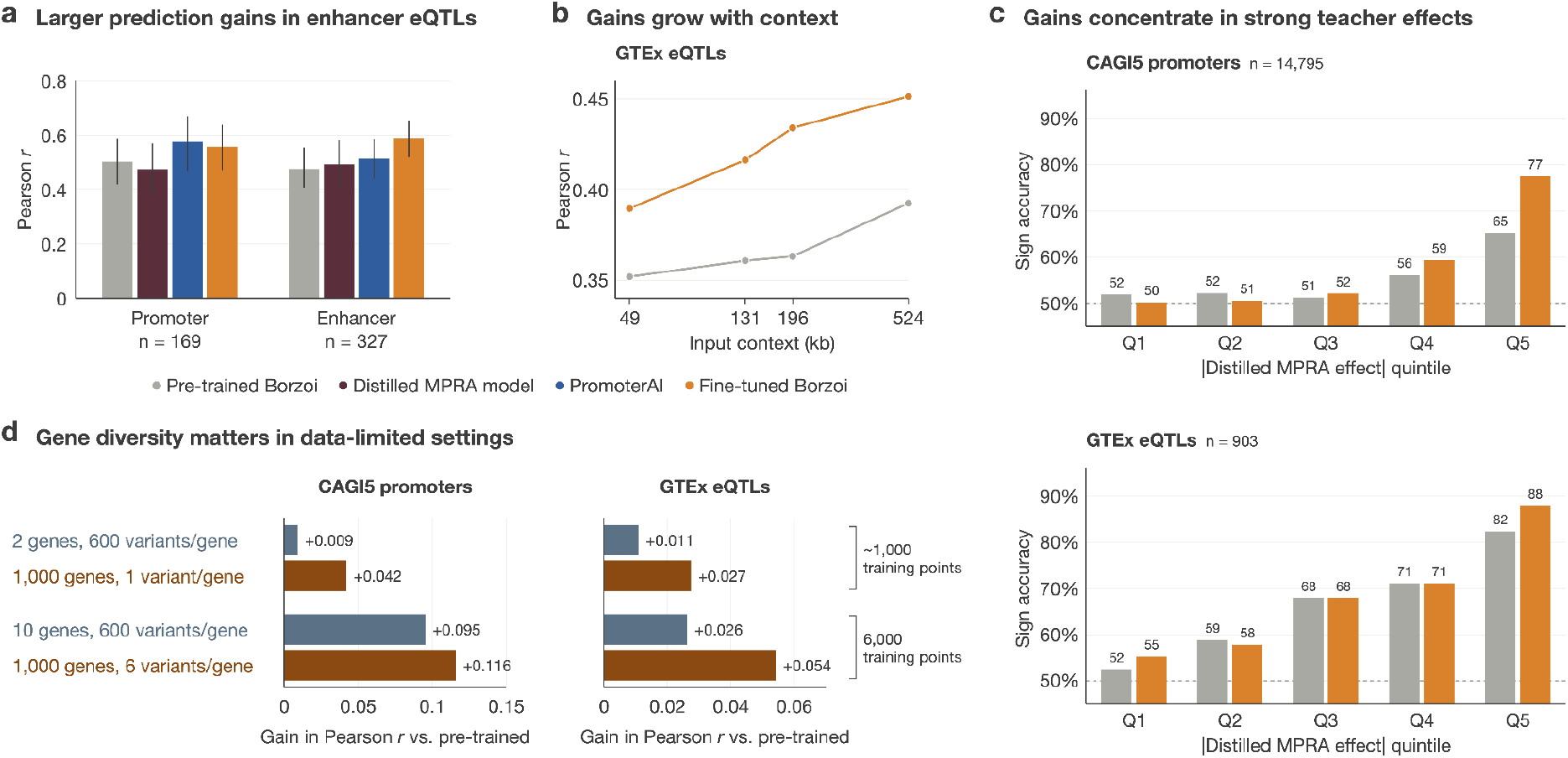
Fine-tuning gains depend on context, scale, and regulatory signal. **a**, GTEx whole-blood SuSiE fine-mapped eQTLs (PIP > 0.9) are stratified by ENCODE candidate *cis*-regulatory element (cCRE) annotations into promoter-like (*n* = 169) and enhancer-like (*n* = 327). Pearson correlation between predicted and observed eQTL effect sizes for pre-trained Borzoi, distilled MPRA model, PromoterAI, and fine-tuned Borzoi. **b**, Effect of genomic input context on variant-effect prediction performance for pre-trained and fine-tuned Borzoi using GTEx whole-blood SuSiE fine-mapped eQTLs (*n* = 903, PIP > 0.9). Models were evaluated using input windows of 49,152, 131,072, 196,608, and 524,288 bp. Performance was measured by Pearson correlation between predicted and observed eQTL effect sizes. **c**, CAGI5 promoter variants *(left)* and SuSiE fine-mapped whole-blood eQTLs with PIP > 0.9 *(right)* were stratified into quintiles according to the absolute magnitude of effect predicted by the distilled MPRA model. Bars show the percentage of variants for which pre-trained or fine-tuned Borzoi correctly predicts the direction of effect. **d**, In data-limited settings, training on more genes with fewer variants per gene performed better than training on fewer genes with more variants per gene, given the same number of total variants. Models were assessed on (left) CAGI5 promoter saturation mutagenesis (*n* = 5,769) and (right) GTEx whole-blood SuSiE fine-mapped eQTLs, PIP > 0.9 (*n* = 903).

We next evaluated whether the fine-tuned Borzoi model could be further improved by increasing the input sequence size at inference time, providing more genomic sequence context for each prediction. We evaluated the same set of fine-mapped GTEx eQTLs across context lengths, expanding the exon output bins to include additional target-gene exons captured by larger windows. Accuracy on GTEx eQTLs increased with input length, rising from Pearson *r* = 0.39 at 49,152 bp to 0.45 at 524,288 bp (Fig. 3b), and exceeded pretrained Borzoi at every length tested. Thus, short-window fine-tuning retained the predictive benefit of longer genomic inputs. We used 524,288-bp windows in all subsequent analyses. To identify which variants benefited most from fine-tuning, we stratified CAGI5 promoter variants and GTEx eQTLs into quintiles by their magnitude of effect predicted by the distilled MPRA model (Fig. 3c). We then measured sign accuracy, the fraction of variants whose predicted effect direction matched the measured MPRA or eQTL effect. Gains in sign accuracy from fine-tuning were concentrated in the top quintile (CAGI5: 65% to 77%; GTEx: 82% to 88%) of effects predicted by the distilled MPRA model. These findings suggest that synthetic MPRA supervision is particularly effective for variants with larger predicted MPRA effects, consistent with previous work showing that S2F models perform best on variants with strong predicted effects^20^.

Finally, we asked how label volume and diversity affected model accuracy. We scaled the training data in two ways: by increasing the number of genes but fixed at 600 variants per gene, and separately by increasing the number of variants per gene with the gene set fixed at 1,000. Prediction accuracy generally improved as training-set size increased for both CAGI5 promoter variants and 903 GTEx whole-blood eQTLs from 824 genes held out from training (Supplementary Fig. 3). Overall, training on more genes with fewer variants per gene yielded greater improvements than training on fewer genes with more variants per gene (Fig. 3d). At approximately 1,000 total training variants, Pearson *r* increased relative to pretrained Borzoi by 0.042 versus 0.009 on CAGI5 and by 0.027 versus 0.011 on GTEx eQTLs from 824 held-out genes. This advantage persisted at 6,000 total training variants, with gains of 0.116 versus 0.095 on CAGI5 and 0.054 versus 0.026 on GTEx eQTLs. The difference between the two scaling strategies diminished at larger training sizes, suggesting that gene diversity matters most when data are limited (Supplementary Fig. 3).

### Fine-tuned Borzoi learns novel regulatory features

To interpret how synthetic MPRA supervision alters the sequence features Borzoi identifies, we computed input × gradient attributions across 195 held-out genes and called seqlets and motifs with Tangermeme^21^ for pre-trained and fine-tuned Borzoi (Fig. 4a). Fine-tuned Borzoi identified 2%, 26%, and 71% more seqlets than pre-trained Borzoi at *p* < 0.05, 0.01, and 0.001, respectively. The gains grow at stricter thresholds, suggesting that fine-tuning sharpens attributions at regulatory features. To test whether these changes reflect information from the distilled MPRA model, we compared motifs identified by all three models within the 281-bp window centered on each gene’s TSS (*p* < 0.01; Fig. 4b). Of the 491 motifs identified by fine-tuned Borzoi, it shared 50.1% of the distilled MPRA model’s 357 motifs and retained 54.0% of pre-trained Borzoi’s 463, while 132 were unique to the fine-tuned model. Fine-tuned Borzoi thus combines regulatory features from both of its source models and also generalizes to additional features. For example, at the promoter of the held-out gene *NUCB1*, both the distilled MPRA model and fine-tuned Borzoi identified a motif matching ZNF143 (*p* = 0.006 and 0.009, respectively), whereas pre-trained Borzoi did not (Fig. 4c). Together, these results suggest that synthetic variant effect supervision teaches Borzoi regulatory features that transfer to unseen genes, while preserving much of its pre-trained regulatory knowledge.

**Figure 4:**
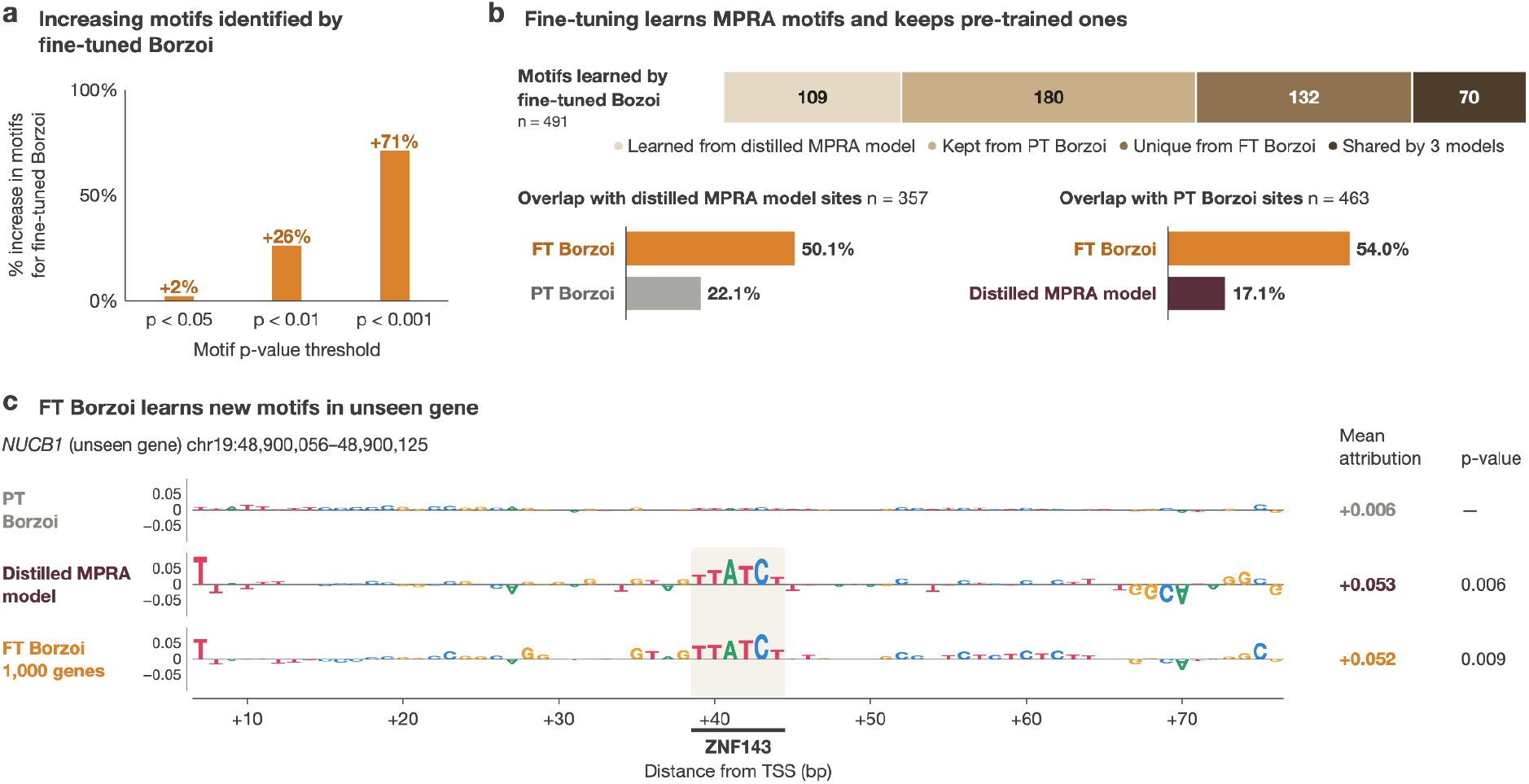
Fine-tuned Borzoi learns generalizable regulatory sequence features. **a**, Number of motif seqlets identified based on fine-tuned and pre-trained Borzoi input x gradient attributions across 195 held-out genes. Seqlets were identified using tangermeme at three significance thresholds (*p* < 0.05, 0.01, and 0.001) and matched against HOCOMOCO v13 motifs. **b**, Overlap of regulatory motifs identified by pre-trained Borzoi, distilled MPRA student, and fine-tuned Borzoi across 195 held-out genes. **c**, Input x gradient attributions for pre-trained Borzoi, the distilled MPRA model, and fine-tuned Borzoi at *NUCB1*, an unseen gene. Mean attribution and *p-*value show significant motif hits in both fine-tuned Borzoi and the MPRA distilled model, but not for the pre-trained Borzoi.

**Supplementary Figure 1:**
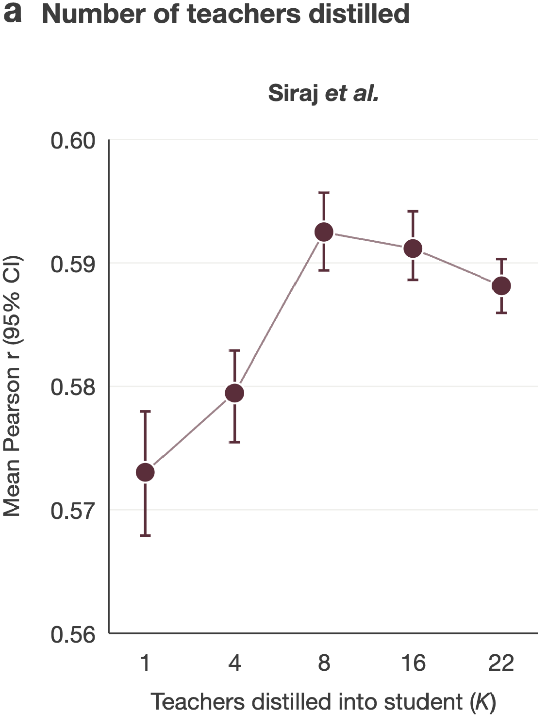
Effect of teacher ensemble size on distilled MPRA model performance. Mean within-element Pearson *r* on Siraj *et al*. K562 MPRA activity for MPRA student models distilled from increasing numbers of teachers (*K* = 1, 4, 8, 16 and 22). Error bars represent 95% confidence intervals.

**Supplementary Figure 2:**
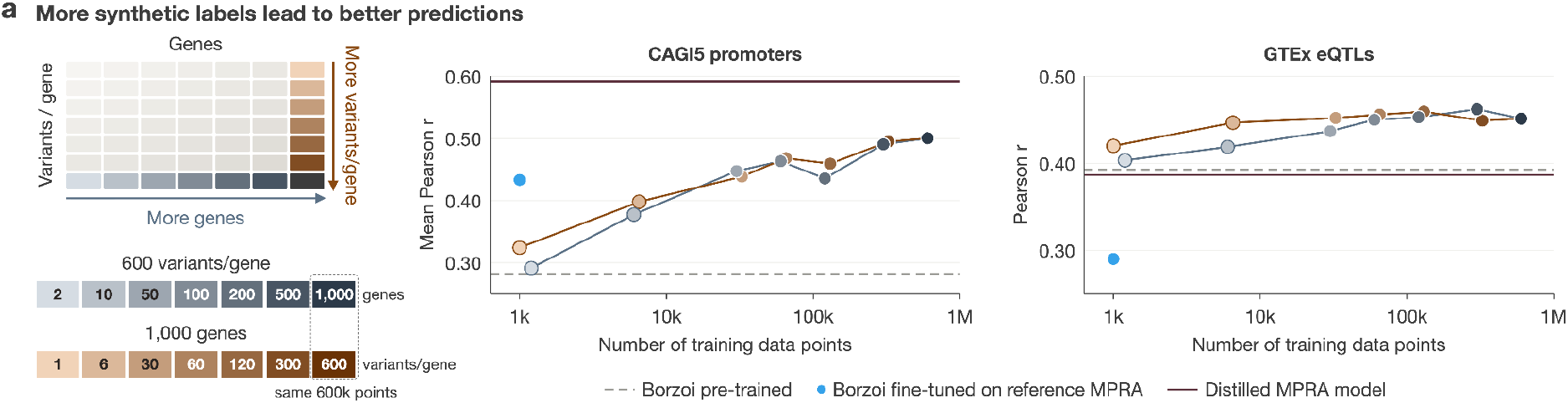
Scaling synthetic supervision across genes and variants. Fine-tuned Borzoi performance with increasing synthetic training labels, either by increasing the number of genes at 600 variants per gene (*blue*) or the number of variants per gene across 1,000 genes (*orange*). The schematic (*left*) illustrates both strategies. Pearson correlation was evaluated on CAGI5 promoter variants (*n* = 5,769) and GTEx Whole Blood fine-mapped eQTLs (*n* = 903, PIP > 0.9).

## Discussion

Synthetic variant effect supervision improved fine-tuned Borzoi’s predictions for unseen variants and genes, including enhancer variants outside the promoter-derived training labels, while preserving pre-trained predictions. Our framework draws large sets of synthetic labels from smaller models trained on existing experimental MPRA data, providing large-scale supervision without new perturbation experiments. Performance also continued to improve as synthetic labels increased. As functional genomics datasets and specialized models continue to expand, this offers a practical way to integrate their regulatory knowledge into large S2F models, given that comprehensive perturbation experiments remain costly.

Our results also suggest ways to make synthetic supervision more efficient. Fine-tuning gains were concentrated among variants with large predicted effects, and gene diversity mattered more than variant density when training data were limited. Focusing supervision on likely regions of large regulatory effect could direct training toward informative variants and allow the same number of training examples to cover more genes and regulatory elements, including enhancers. Such regions could be defined by accessible chromatin from DNase- or ATAC-seq, or by motifs identified by the MPRA model.

Label quality could also be improved: although ensemble distillation reduces noise in synthetic labels^1,17^, weighting labels by estimated reliability, such as teacher uncertainty^22^, could limit the influence of unreliable predictions. Finally, extending our framework to models trained on complementary assays could improve predictions across regulatory mechanisms and cellular contexts. More broadly, our findings show that existing models can be repurposed to generate synthetic data and overcome experimental data limitations, offering a scalable path toward more accurate and generalizable variant effect prediction.

## Data availability

The CAGI5 data can be found at https://www.nature.com/articles/s41467-019-11526-w. The GTEx_v8 eQTL data can be found at gtexportal.org. The Siraj et al. K562 dataset can be found at https://www.nature.com/articles/s41586-026-10121-6. The eQTLGen Phase I dataset can be found at https://www.nature.com/articles/s41588-021-00913-z.

## Author contributions

L.W. project conceptualization, formal analysis, investigation, methodology, data visualization, and writing. J.G. project conceptualization, methodology, and writing. E.Z. project conceptualization, data visualization, and writing. R.K. data visualization and writing. A.B. project conceptualization, methodology, funding acquisition, and writing.

## Acknowledgements

We’d like to thank members of the Battle Lab for helpful discussions related to this work. This work was funded by R35GM139580 awarded to A.B.

## Methods

### MPRA Dataset

We trained MPRA teachers on the K562 lentiMPRA measurements from Agarwal et al. (2025). We retained 110,359 autosomal elements: 83,481 enhancers, 14,417 promoters, 12,223 elements across six loci centered on human disease-associated and erythroid lineage genes, and 238 positive and negative genomic controls. Each element consisted of a 281-bp MPRA construct, with 200-bp genomic sequence, 15-bp upstream and downstream adapters, 15-bp barcode, and 36-bp minimal promoter. Measured element activity was log_2_(RNA/DNA) averaged across 3 replicates.

### GTEx and eQTLGen benchmark datasets

We restricted *cis*-eQTLs from GTEx v8^15^ whole blood to those variants fine-mapped by SuSiE with posterior inclusion probability exceeding 0.9 (PIP > 0.9). To assess generalization to genes unseen during training, we excluded variants whose eGene was among the 1,000 training genes used for model distillation and fine-tuning. We further required each eGene to have an annotated exon (GENCODE v.26) within the Borzoi input window centered at the variant to calculate gene-level exonic sum, resulting in 903 variant-gene pairs. We constructed 524,288-bp reference and alternative sequences centered on the variant and compared our predictions to each variant’s marginal effect size estimated in GTEx from their eQTL summary statistics.

We additionally evaluated our models on *cis*-eQTL effect sizes from the eQTLGen Phase I dataset^23^, retaining associations with FDR < 0.05. Per gene, we retained the variant with the largest magnitude of effect and excluded genes that were included during fine-tuning. We yielded 11,684 variant–gene pairs, of which 11,607 could be evaluated using 524,288-bp reference and alternative sequences centered on each variant. We further restricted variants to those that are within 10 kb of the gene’s transcription start site (annotated by GENCODE v.26), resulting in 3,539 variant–gene pairs (830 within 1 kb and 2,709 between 1 and 10 kb).

### CAGI5 saturation mutagenesis

The CAGI5 regulatory challenge data^24^ include saturation mutagenesis MPRA measurements of 29 promoter and enhancer elements assayed in 12 cell types. MPRA teacher and distilled models were evaluated on all variants of the 29 CAGI5 element datasets (*n* = 39,170 variants). Fine-tuned Borzoi was evaluated on the CAGI5 promoter subset defined by the PromoterAI evaluation protocol, which excludes variants with SpliceAI score > 0.05 and variants that may overlap with GTEx rare variants. This subset includes *GP1BB, HBB, HBG1, HNF4A, MSMB, LDLR, PKLR*, and *TERT* across 5,769 variants.

### Siraj *et al*. K562 saturation mutagenesis

We used two MPRA datasets from Siraj et al.^16^ to evaluate variant-effect prediction accuracy. To evaluate the MPRA teacher and distilled student models, we used the saturation-mutagenesis MPRA dataset, comprising approximately 200-bp regulatory elements assayed in K562 cells. We excluded indels, multiallelic variants, and elements overlapping the Agarwal et al. lentiMPRA training library, retaining 102 elements (*n* = 150,548 variants). Prediction accuracy was evaluated using Pearson correlation between predicted and experimentally measured log2 allelic skew within each element, averaged across elements.

To evaluate pre-trained and fine-tuned Borzoi, we used naturally occurring expression-modulating variants (emVars) assayed by Siraj et al. in K562 cells. We excluded variants overlapping Agarwal et al. training elements, variants used for PromoterAI training, and variants that overlapped with the 1,000 Borzoi fine-tuning genes, retaining 14,172 emVars. We computed the Pearson correlation between predicted variant effects and experimentally measured log_2_ allelic skew across all retained variants.

### Gene selection and data splits

For all 18,380 autosomal protein-coding genes in GTEx v8 whole blood, we fit an elastic net regression predicting log(TPM + 1) from personal sequences of 670 individuals in a 200 kb window centered on the transcription start site (TSS), as annotated by GENCODE v.26. Per-gene genotype matrices were built with hottools where only biallelic SNVs were retained and encoded in dosages of 0, 1, or 2. We estimated *cis*-heritability as the *R*^*2*^ between the predicted and observed expression in held-out individuals. Of the 1,436 genes with *R*^*2*^ > 0.01, 1,202 overlapped with genes included in the Agarwal et al. dataset. These 1,202 genes were split into 1,000 training genes, 98 validation genes, and 104 test genes for downstream model training and evaluation. Training genes had *R*^*2*^ of 0.01–0.78.

### MPRA teacher training

We fine-tuned the pre-trained AlphaGenome^1^ encoder to predict lentiMPRA activity^13^ following the framework described by Murphy *et al*^*14*^. We used the same 281-bp construct as input, MPRA-head architecture, data augmentation strategy, and two-stage fine-tuning procedure described in their framework. The 110,359 autosomal elements were partitioned by chromosome into 16 folds, and one teacher was trained per fold, yielding 16 MPRA teachers.

### Ensemble distillation of MPRA teachers

The 16 teacher models were distilled into a single MPRA student model with identical architecture and 281-bp input. Distillation data were restricted to enhancer and promoter elements that mapped to the 1,000 training genes. Using *in silico* mutagenesis, we generated all possible single-nucleotide substitutions at every position of each 200-bp element. Each teacher independently predicted the MPRA activity for both the reference and alternate alleles with the variant at four different positions (the center position and three positional offsets that are +/-20 bp from the center) in both forward and reverse-complement directions. These predictions were averaged within each teacher and subsequently across 16 teachers to obtain ensemble-level reference (*y*_*ref*_) and alternate predictions (y_alt_). The MPRA student model *f(x)* was trained to reproduce the ensemble-averaged predictions for the reference and alternate sequences by minimizing the loss *L* = MSE(*f(x*_*ref*_*), y*_*ref*_) + MSE(*f(x*_*alt*_*), y*_*alt*_). The reference-only distilled student (Fig. 2b) was trained identically but with the loss function only minimizing MSE(*f(x*_*ref*_*), y*_*ref*_). To assess the effect of ensemble size (Supplementary Fig. 1), students were also distilled from 1, 4, 8, 16, and 22 teachers with the same distillation data and training procedure as the 16-teacher student.

### Fine-tuning Borzoi on synthetic variant effect predictions

We fine-tuned Borzoi (replicate 0) on synthetic variant effect predictions for all 600 SNVs (three alternative alleles at each of the 200 positions) in the 200-bp core promoter element for each one of 1,000 training genes (total training = 600,000 variants). Whereas the distilled MPRA student scores a 281-bp construct, Borzoi received each variant in a genomic context of 49,152-bp, with the variant at the center for both reference and alternative sequences.

The synthetic label of variant *i* is defined as *y*_*i*_ = *f(x*_*alt,i*_) – *f(x*_*ref,i*_), which is the difference between the distilled MPRA student’s predicted alternative- and reference-allele activity, each averaged over the four positions and two orientations (same as distillation). Borzoi’s predicted variant effect size was computed from the three GTEx whole-blood RNA-seq outputs (tracks 7,531–7,533). For each sequence input, predicted coverage was summed over the 32-bp output bins overlapping any exon of the target gene (annotated by GENCODE v.26) and averaged across the three RNA-seq tracks, giving *S*_*ref*_ and *S*_*alt*_ for reference and alternative sequence predictions. The predicted variant effect of Borzoi is *y = log*_*2*_*(S*_*alt*_ *+ S*_*ref*_).

Borzoi was fine-tuned using a pairwise loss over *B* variants per batch, 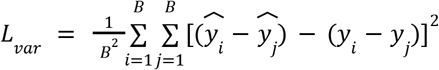 , where *i* and *j* are index variants and *B* = 16. The pairwise loss matches relative differences in effect sizes between variants. To keep reference-sequence predictions close to those of pre-trained Borzoi, we applied a Poisson loss^3^ between the fine-tuned model’s predictions (s) and pre-trained Borzoi’s predictions (t) on the reference sequence across all 7,611 output tracks: 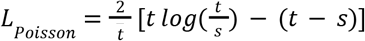 , where 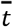 is the mean pre-trained predicted coverage. The total loss was *L* = *L*_*Pairwise*_ + 0.5 *L*_*Poisson*_.

### Variant scoring

Unless otherwise stated, pre-trained and fine-tuned Borzoi scored each variant with matched 524,288-bp reference and alternative sequences centered on the variant, summing over exons of the eGene or gene of the CAGI5 promoter element. Exons were annotated by GENCODE v.26. MPRA models scored each variant as the difference between predicted alternative- and reference-allele activity of 281-bp constructs centered on the variant. MPRA-LegNet, PARM, and PromoterAI were also scored similarly, each with the input length on which it was trained (230 bp, 600 bp, and 20,480 bp, respectively).

### Scaling of synthetic supervision

We fine-tuned Borzoi on training sets scaled in two ways: increasing the number of genes (2, 10, 50, 100, 200, 500, or 1,000 genes, each with all 600 variants) or increasing the number of variants per gene (1, 6, 30, 120, 300, or 600 variants for each of the 1,000 training genes). Both scaling methods end with the same 600,000-variant training set. As a control, Borzoi was also fine-tuned on the promoter MPRA activity corresponding to the 1,000 genes from the Agarwal *et al*. dataset.

### Attribution and motif analysis

For 195 of the 202 held-out genes for which a full 49,152-bp input window centered on the TSS could be constructed, we computed input x gradient attributions of pre-trained and fine-tuned Borzoi on 49,152-bp sequences centered on the TSS. Because the distilled MPRA model accepts only 281-bp inputs, motifs were compared across the three models on attributions computed from the same 281-bp sequence centered on each TSS. Seqlets were identified with tangermeme^21^ at *p* < 0.05, 0.01, and 0.001 and annotated with HOCOMOCO v.13^25^ transcription factor motifs, retaining matches with *q* < 0.05.

